# Brief online mindfulness meditation training modulates heart rate variability during post-stress meditation

**DOI:** 10.64898/2026.02.11.705451

**Authors:** Yuki Tsuji, Itsuki Kondo, Sotaro Shimada

**Author notes:** Corresponding author: Yuki Tsuji ( /), Meiji University, 1-1-1 Higashimita, Tama-ku, Kawasaki-city, Kanagawa, 214-8571, Japan.

## Abstract

Mindfulness-based interventions are increasingly delivered online, yet evidence for short programs often relies on self-report outcomes. We tested whether a brief online mindfulness meditation training produces detectable changes in autonomic regulation during a standardized stress-to-meditation sequence. Healthy adults with no meditation experience were randomized to a four-week online mindfulness meditation program (MG) or an active health-management program (CG). Before and after training, participants completed a laboratory session consisting of rest, a mental arithmetic stress task, guided focused-attention breathing meditation, and post-rest while ECG was recorded. Across the training period, both groups showed reduced negative affective symptoms, but only the mindfulness group showed an increase in the Observing facet. Critically, frequency-domain HRV indices during the laboratory protocol showed a group-specific post-training pattern: MG exhibited lower LF/HF and higher normalized HF power (nHF) compared with pre-training, and MG differed from CG in the post-training session. Within MG, training-related improvement in FFMQ Non-reactivity was positively associated with nHF during the post-stress meditation period. These findings indicate that a brief online mindfulness program can modulate HRV during a stress-to-meditation context and that post-stress autonomic modulation during meditation covaries with acceptance-related skill acquisition.

## Introduction

Mindfulness-based interventions (MBIs) are widely used to reduce stress-related symptoms and to cultivate attentional and acceptance-related skills. Mindfulness is often described as present-moment awareness cultivated through purposeful, non-judgmental attention to experience^[1]^. Although brief online mindfulness programs have expanded rapidly, much of the evidence for short interventions relies primarily on self-report outcomes, leaving open whether such programs yield detectable changes in objective indices of stress-related physiology. Meta-analytic evidence indicates that online MBIs can produce small-to-moderate improvements in mental health outcomes such as anxiety, depression, and stress, alongside increases in self-reported mindfulness^[2–4]^. With the rapid expansion of remote delivery formats, brief online mindfulness programs have become a practical option for reaching large populations with low barriers to access, highlighting the need for objective physiological evidence for short programs^[5–7]^.

Heart rate variability (HRV) provides a noninvasive index of autonomic regulation and has been widely used as a psychophysiological marker in stress and emotion research^[8,9]^. Prior work suggests that mindfulness training and meditation practice can be accompanied by increases in vagally mediated HRV indices, consistent with enhanced parasympathetic cardiac modulation. HRV has also been examined in cognitively demanding stress contexts; for example, a mindfulness training study in teacher trainees reported higher HRV during a cognitive challenge relative to control conditions^[10]^. Moreover, brief online interventions have begun to incorporate HRV in diverse settings (e.g., acute stress manipulations or naturalistic monitoring)^[11,12]^. Together, these lines of evidence support HRV as a plausible objective outcome for evaluating remotely delivered mindfulness training.

One underexplored context is guided meditation immediately following acute cognitive stress within a standardized session. This post-stress meditation sequence is practically relevant to real-world stress regulation, because brief mindfulness practice can be used as an immediate strategy to down-regulate stress after demanding events. Yet, few studies have directly evaluated whether a short online mindfulness program produces group-specific changes in autonomic indices measured under a protocol that combines an acute cognitive stressor with subsequent focused-attention (FA) breathing meditation, while also testing whether physiological changes covary with training-related changes in mindfulness skills. Mechanistic accounts emphasize that mindfulness training involves both monitoring/attentional stability and acceptance-related processes, and that acceptance-oriented skills can be central for translating monitoring into stress-relevant benefits^[13]^. The Five Facet Mindfulness Questionnaire (FFMQ) is a widely used measure of dispositional mindfulness that allows these skills to be assessed at the facet level^[14]^, providing a useful framework for linking training-related changes in subjective skills to physiological modulation during practice.

The present study aimed to clarify the effects of a brief, four-week online mindfulness meditation training program using both subjective self-report measures and HRV indices. Participants were randomly assigned to a mindfulness meditation training group (MG) or a health-management control group (CG) and completed weekly self-report assessments during the four-week period. They also underwent pre- and post-training laboratory sessions that included an acute cognitive stress task followed by an audio-guided FA breathing meditation. A health-management program served as an active comparison condition. We tested whether the mindfulness program would be associated with (i) changes in mindfulness-related self-report facets and (ii) group-specific shifts in HRV indices during the standardized post-stress meditation protocol. We further examined whether individual differences in training-related improvement in mindfulness skills—particularly acceptance-related facets—would covary with HRV during the post-training meditation period, providing convergent evidence linking skill acquisition to physiological modulation.

## Results

This study evaluated a brief, four-week online mindfulness meditation training program using self-report measures collected across the training period and ECG-derived HRV indices obtained during pre- and post-training laboratory sessions. The experiment consisted of three phases: pre-training, training, and post-training. In both laboratory sessions (pre- and post-training), participants completed a standardized four-period protocol comprising a pre-rest period, an acute cognitive stress task (mental arithmetic), an audio-guided focused-attention (FA) breathing meditation period, and a post-rest period, with ECG recorded during each period. During the training phase, outcomes were compared between a mindfulness meditation training group (MG) and an active health-management control group (CG) (Figure 1). The MG participated in a four-week online meditation program with daily practice, with the meditation technique varying by week. The CG participated in a four-week health-management program that included nutrient intake intended to reduce mental distress and a food diary, with the weekly theme varying by week. A weekly meeting was held to review the previous week and outline the next week’s theme. Participants completed questionnaires assessing mindfulness (Five Facet Mindfulness Questionnaire; FFMQ), depressive symptoms (Beck Depression Inventory–II; BDI-II), and anxiety (State–Trait Anxiety Inventory; STAI) before the pre-training session and each weekend during the training phase.

**Figure 1.**
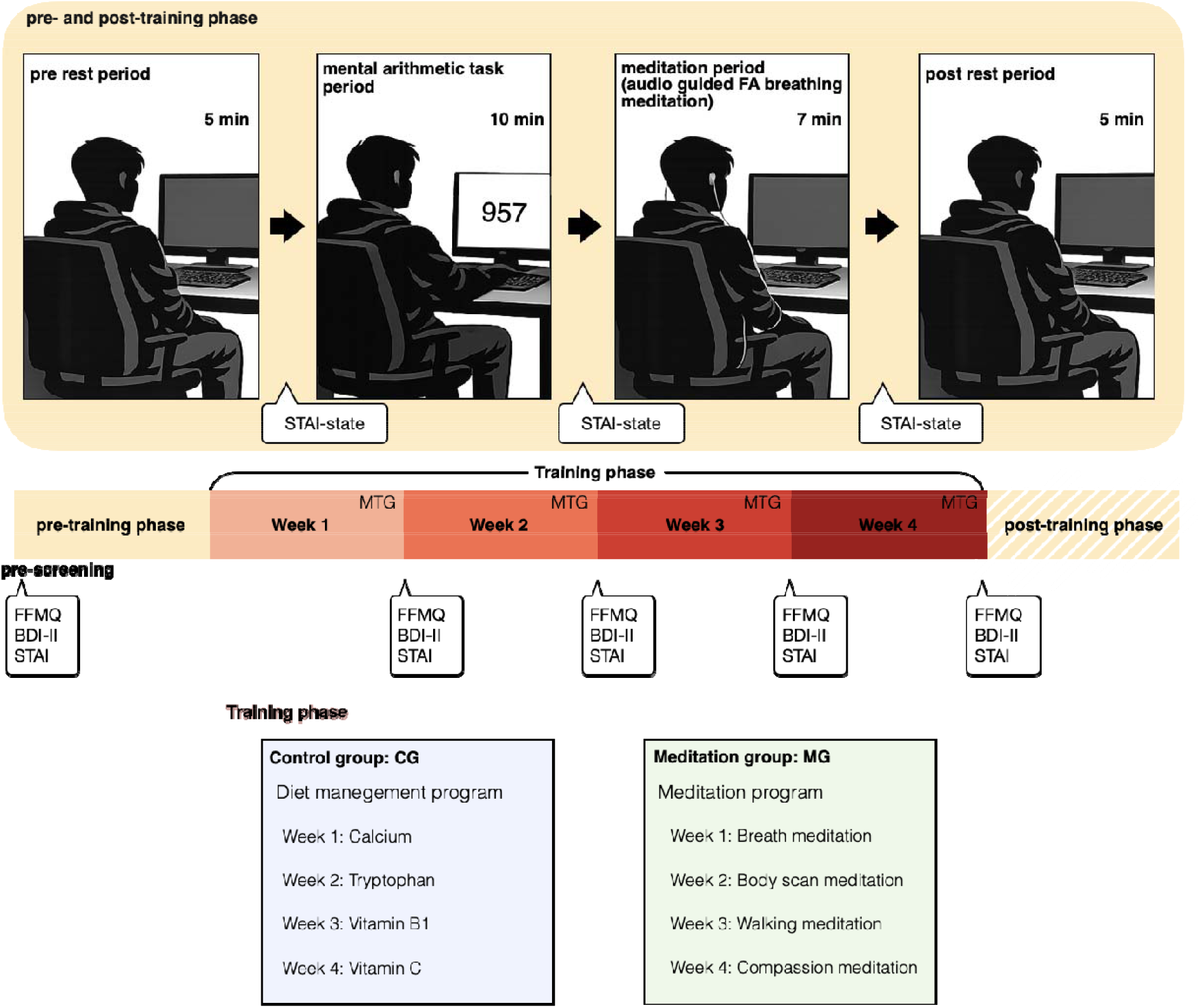
Experimental procedure.

### Psychological scales

At pre-screening, no significant differences were observed between groups in the results of the psychological scales (FFMQ, STAI, and BDI-II) (see Table S1).

Week-to-week changes in these self-report measures across the training period are illustrated in Figure 2. To characterize week-to-week changes in self-report measures across the training period, we conducted two-way ANOVAs (Week × Group) with aligned rank transform (ART)^[15]^ for each questionnaire outcome, followed by Wilcoxon signed-rank tests with Holm correction when a significant main effect of Week was detected. The *F* statistics and effect sizes for the Week main effect, together with the outcomes of the corrected post hoc comparisons, are summarized in Table 1. For FFMQ, FFMQ describing, and STAI-trait, the two-way ANOVA revealed the significant main effect of Week (FFMQ: *F*(4, 152) = 2.65, *p* < 0.05, *η*_*p*_^2^ = 0.0652, FFMQ describing: *F*(4, 152) = 2.59, *p* < 0.05, *η*_*p*_^2^ = 0.0638, STAI-trait: *F*(4, 152) = 4.22, *p* < 0.05, *η*_*p*_^2^ = 0.100). Post hoc analysis (the Wilcoxon’s signed-rank test with Holm’s p-value adjustment) for the significant main effect of Week revealed that there was no difference between weeks in the scores of the FFMQ total, describing, STAI-trait. For BDI-II, the two-way ANOVA revealed the significant main effect of Week (*F*(4, 152) = 8.60, *p* < 0.01, *η*_*p*_^2^ = 0.185). The Wilcoxon’s signed-rank test was performed and corrected for multiple comparisons with Holm’s p-value adjustment. The result showed that the BDI-II score at pre-screening was higher than that observed at week 3 (*Z* = 547, *p* < 0.01, *r* = 0.554) and week 4 (*Z* = 553, *p* < 0.05, *r* = 0.493), and the BDI-II score at week 2 was higher than that observed at week 3 (*Z* = 406, *p* < 0.01, *r* = 0.510) and week 4 (*Z* = 461, *p* < 0.05, *r* = 0.530). In addition, the BDI-II score at week 1 was higher than that observed at week 3 (*Z* = 487, *p* < 0.01, *r* = 0.456) (Figure 2 A). For STAI-state, the two-way ANOVA revealed the significant main effect of Week (*F*(4, 152) = 4.04, *p* < 0.01, *η*_*p*_^2^ = 0.0962). The Wilcoxon’s signed-rank test was performed and corrected for multiple comparisons with Holm’s p-value adjustment.

**Table 1.** Week effects on psychological scale scores during the training period. Two-way ANOVAs (Week × Group) were conducted for each scale; the *F* statistics and partial eta-squared (*η*_*p*_^2^) for the main effect of Week are shown, along with Holm-corrected post hoc comparisons (Wilcoxon signed-rank tests). (n.s., no significant pairwise differences after Holm correction; \**p* < 0.05, \*\**p* < 0.01.).

| Psychological scales | Week main effect | Post hoc comparisons |
| --- | --- | --- |
| FFMQ |  |  |
| total | $F(4,152) = 2.65^*$, $\eta_p^2 = 0.0652$ | n.s. |
| describing | $F(4,152) = 2.59^*$, $\eta_p^2 = 0.0638$ | n.s. |
| BDI-II | $F(4,152) = 8.60^*$, $\eta_p^2 = 0.185$ | Pre-screening > Week3**<br>Pre-screening > Week4**<br>Week1 > Week3*<br>Week2 > Week3**<br>Week2 > Week4** |
| STAI |  |  |
| state | $F(4,152) = 4.04^*$, $\eta_p^2 = 0.0962$ | Pre-screening > Week3**<br>Pre-screening > Week4**<br>Week2 > Week4** |
| trait | $F(4,152) = 4.22^*$, $\eta_p^2 = 0.100$ | n.s. |

**Figure 2.**
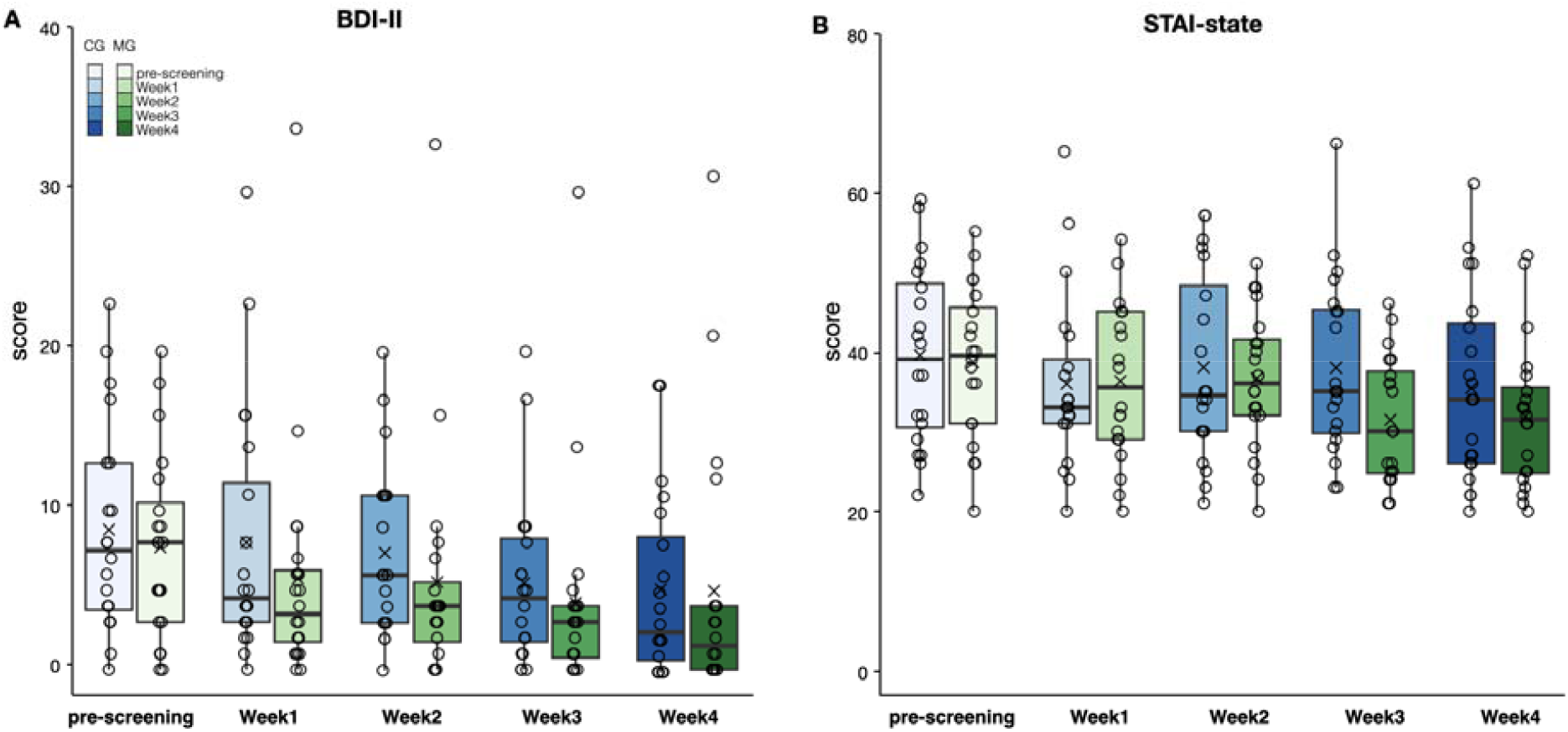
The box plots illustrate the median and interquartile range of the BDI-II (**A**) and STAI-state (**B**) score at each week (pre-screening, week 1, week 2, week 3, and week 4). The mean values of each week are represented as crosses and individual data points are represented as circles.

The result showed that the STAI-state score at pre-screening was higher than that observed at week 3 (*Z* = 605, *p* < 0.01, *r* = 0.487) and week 4 (*Z* = 494, *p* < 0.05, *r* = 0.553), and the STAI-state score at week 2 was higher than that observed at week 4 (*Z* = 572, *p* < 0.01, *r* = 0.473) (Figure 2 B). No other significant main effect or interaction were observed (Table S2).

State anxiety increased after the stress task and decreased thereafter, with a similar within-session pattern pre/post training and no clear group-specific modulation. To evaluate the effect of training on the STAI state in each period, a three-way ANOVA with ART was performed with Group (MG or CG) as the between-subjects factor, Phase (pre-training or post-training) and Period (after pre-Rest, after stress task or after post-Rest) as the within-subject factors. The three-way ANOVA revealed significant main effects (Phase: *F*(1, 38) = 19.7, *p* < 0.01, *η*_*p*_^2^ = 0.341, Period: *F*(2, 76) = 70.6, *p* < 0.01, *η*_*p*_^2^ = 0.650) and the interaction between Phase and Period (*F*(2, 76) = 19.4, *p* < 0.01, *η*_*p*_^2^ = 0.338).

Post-hoc analysis (simple main effect tests using Friedman’s test or Wilcoxon’s rank test) for interactions revealed that significant main effects of Period (pre-training: χ^*2*^(2) = 52.8, *p* < 0.01, post-training: χ^*2*^(2) = 23.5, *p* < 0.01). To examine differences between periods, the Wilcoxon’s signed-rank test was performed and corrected for multiple comparisons with Holm’s p-value adjustment. At pre-training phase, the STAI state score after stress period task was larger than after pre-Rest period (*Z* = 777, *p* < 0.01, *r* = 0.751) and after post Rest period (*Z* = 10.0, *p* < 0.01, *r* = 0.728). At post-training phase, the STAI state score after stress period task was larger than after pre-Rest period (*Z* = 114, *p* < 0.01, *r* = 0.383) and after post-Rest period (*Z* = 788, *p* < 0.01, *r* = 0.654), and after pre-Rest period larger than after post-Rest period (*Z* = 516, *p* < 0.01, *r* = 0.281). Additionally, the STAI state score after stress task period at pre-training phase was larger than post-training phase (*Z* = 687, *p* < 0.01, *r* = 0.702) and after meditation period at pre-training phase was larger than post-training phase (*Z* = 471, *p* < 0.01, *r* = 0.410). However, no significant main effect of Group or interaction between Group and other factors was observed.

Psychological symptoms improved over time in both groups, whereas the FFMQ observing score increased selectively in the meditation group. To evaluate the effect of training on score of psychological scales, a two-way analysis of variance (ANOVA) with ART was conducted with Group (Meditation group: MG or Control group: CG) as a between-subject factor and Week (pre-screening, week 1, week 2, week 3, or week 4) as a within-subject factor. For FFMQ observing, the two-way ANOVA revealed significant interaction between Group and Week (*F*(4, 152) = 3.33, *p* < 0.05, *η*_*p*_^2^ = 0.0805). Post-hoc analysis (simple main effect tests using Friedman’s test or Wilcoxon’s rank test) for interactions revealed that significant main effect of Week in the MG was observed (χ^*2*^ (4) = 13.9, *p* < 0.01). To examine differences between weeks, the Wilcoxon’s signed-rank test was performed and corrected for multiple comparisons with Holm’s p-value adjustment. The FFMQ observing score at week 4 was higher than that observed at week 1 (*Z* = 18.5, *p* < 0.05, *r* = 0.732) and week 3 showed significantly higher scores than week 1 in the MG (*Z* = 22.5, *p* < 0.05, *r* = 0.687) (Figure 3).

**Figure 3.**
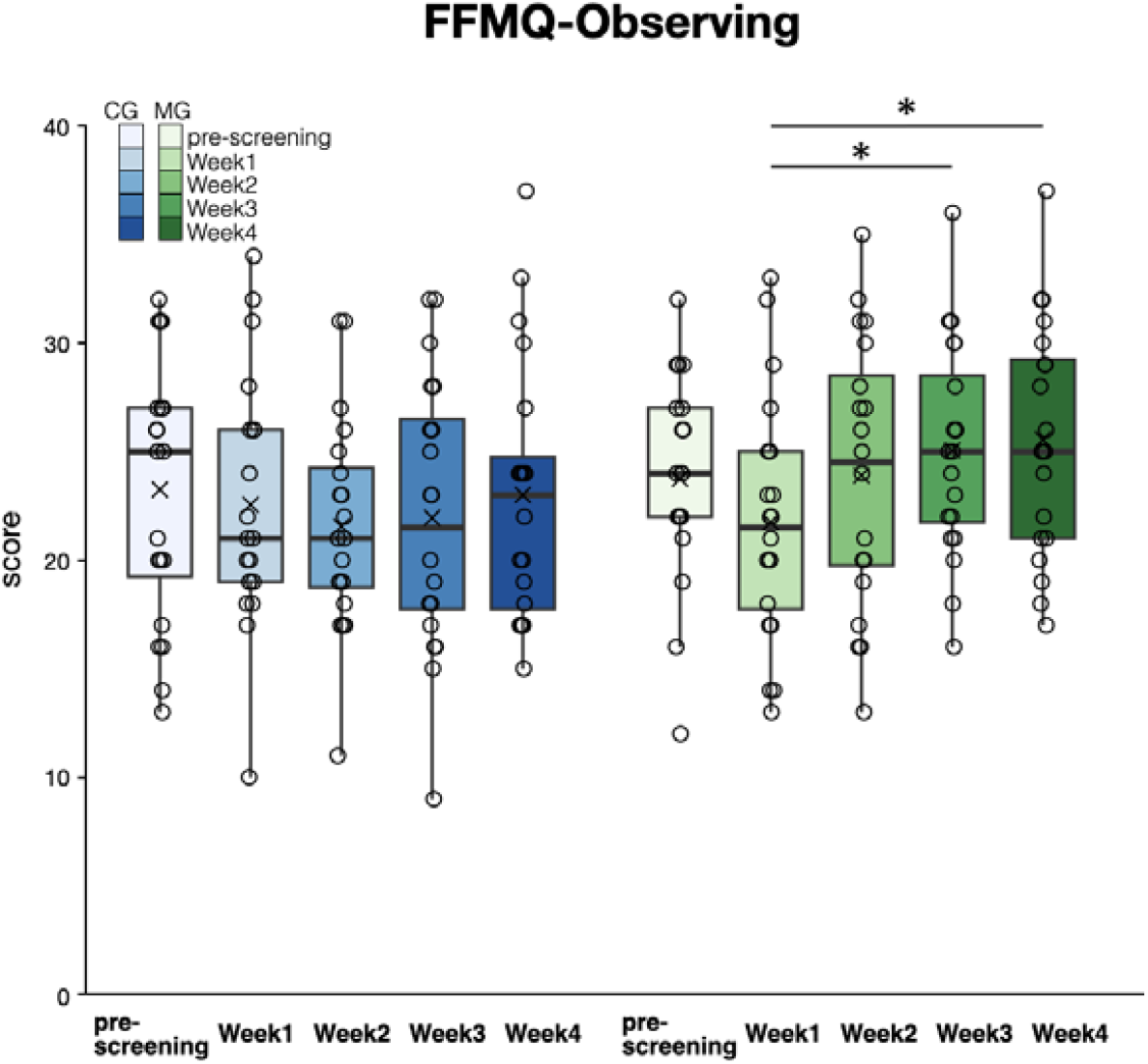
The box plots illustrate the median and interquartile range of the FFMQ observing score at each week (pre-screening, week 1, week 2, week 3, and week 4). The mean values of each week are represented as crosses and individual data points are represented as circles. (*: *p* < 0.05).

### Heart rate variability

In contrast to self-report symptom changes, HRV indices showed a meditation-specific post-training shift during the post-stress meditation, converging in higher nHF and lower LF/HF. To assess autonomic system activities, Heart rate (HR) variability each period was performed frequency analysis. The power ratio of the low-frequency (LF) (0.04 to 0.15 Hz) to high-frequency (HF) (0.15 to 0.4 Hz) band in the HR variability spectrogram (LF/HF) is considered as an index of sympathetic activity, and normalized HF power (nHF = HF/(LF+HF)) is considered as an index of parasympathetic activity. The power ratio of the resting period to other periods (mental arithmetic task, meditation, and post-rest period) was calculated.

To evaluate the effect of training on the LF/HF and nHF, a three-way ANOVA with ART was performed with Group (MG or CG) as the between-subjects factor, Phase (pre-training or post-training) and Period (after pre-Rest, after stress task or after post-Rest) as the within-subject factors. For LF/HF, the three-way ANOVA revealed significant main effects (Phase: *F*(1, 38) = 11.2, *p* < 0.01, *η*_*p*_^2^ = 0.228, Period: *F*(2, 76) = 6.84, *p* < 0.01, *η*_*p*_^2^ = 0.152) and the interaction between Phase and Group (*F*(1, 38) = 12.8, *p* < 0.01, *η*_*p*_^2^ = 0.252). To examine the differences between groups and phases, the Wilcoxon’s signed-rank test was performed and corrected for multiple comparisons with Holm’s p-value adjustment. The results showed that the LF/HF in the post-training phase was significantly lower than that observed in the pre-training phase in the MG (*Z* = 1662, *p* < 0.01, *r* = 0.502). Moreover, the MG exhibited a lower LF/HF than the CG in the post-training phase (*Z* = 2613, *p* < 0.01, *r* = 0.390) (Figure 4 A). For nHF, the three-way ANOVA revealed the significant interaction between Phase and Group (*F*(1, 38) = 14.0, *p* < 0.01, *η*_*p*_^2^ = 0.269). The results of multiple comparisons with Holm’s p-value adjustment showed that the nHF in the post-training phase was significantly greater than that observed in the pre-training phase in the meditation group (*Z* = 387, *p* < 0.01, *r* = 0.355). Moreover, the meditation group exhibited a greater nHF than the control group in the post-training phase (*Z* = 1035, *p* < 0.01, *r* = 0.367) (Figure 4 B).

**Figure 4.**
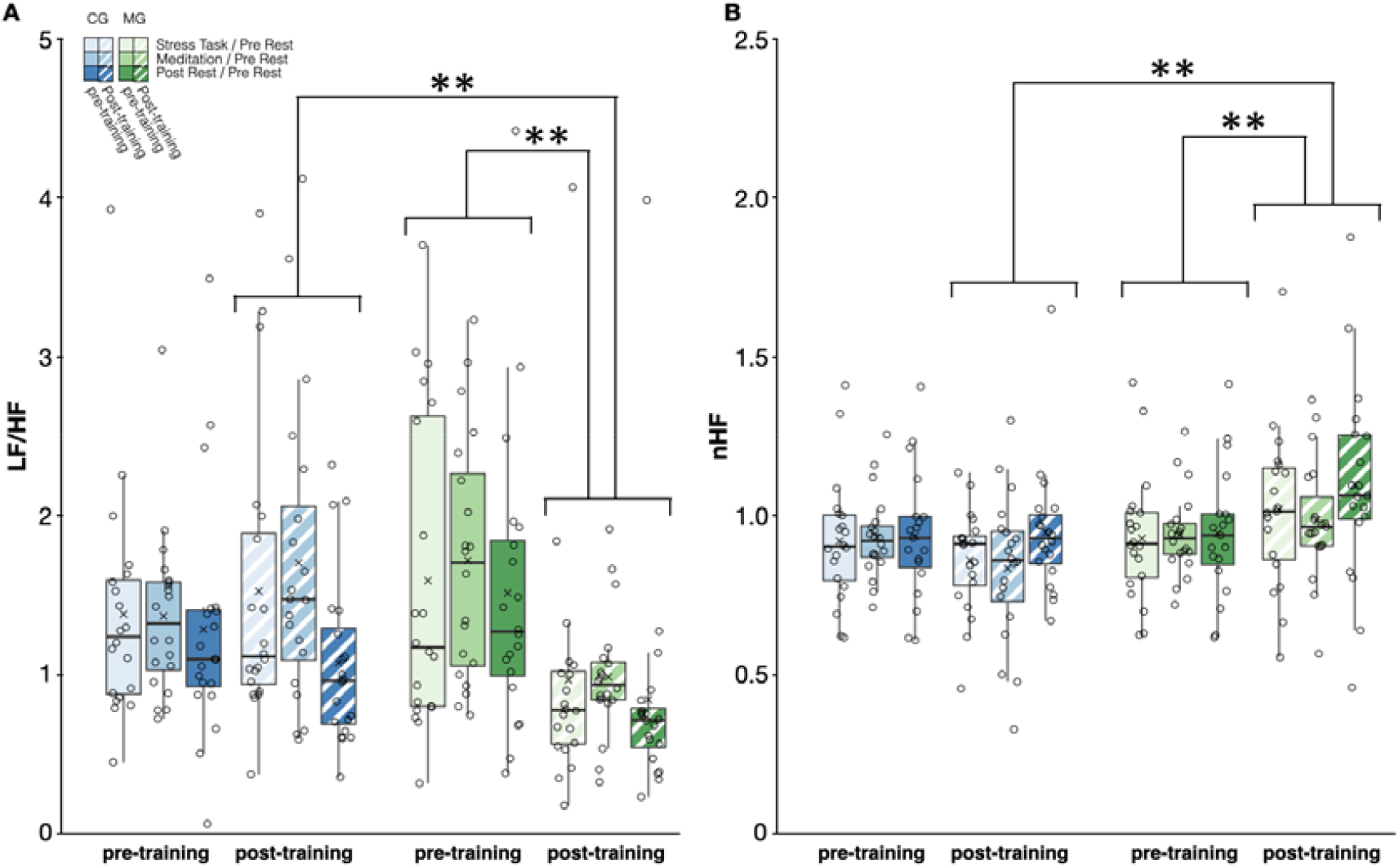
The box plots illustrate the median and interquartile range of the ratio of LF/HF (**A**) or nHF (**B**) for each period to the pre-rest period. Group differences emerged primarily in the post-training session, where the meditation group showed lower LF/HF and higher nHF than the control group. The mean values of each period are represented as crosses and individual data points are represented as circles. (**: *p* < 0.01).

Finally, within the meditation group, individual differences in acceptance-related skill gain (Non-reactivity) tracked parasympathetic modulation during the post-training meditation period. To examine the relationship between autonomic nervous system activities and score of psychological scales, Spearman’s rank correlation coefficients were calculated between LF/HF or nHF during each period (mental arithmetic task, meditation, and post-rest period) in the post-training phase and the difference between psychological scale scores in weeks 1 and 4. The result showed a positive correlation between FFMQ non-reactivity and nHF in the MG during the meditation period (*r*_*s*_ = 0.713, Bonferroni corrected: *p* < 0.05/27 (3 periods × 9 psychological scales), Figure 5). No other significant correlation between autonomic nervous system activities and score of psychological scales was observed.

**Figure 5.**
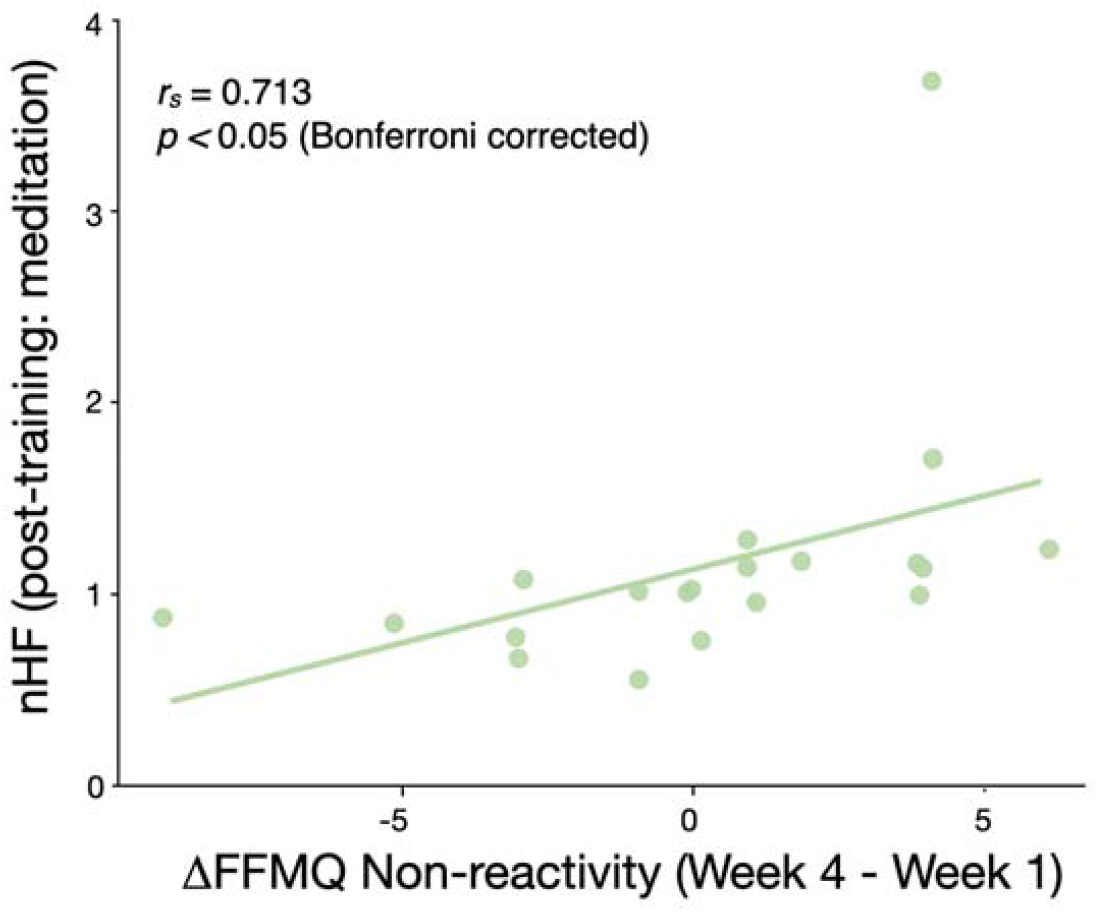
Correlation between FFMQ Non-reactivity gain and nHF during post-stress meditation in the MG. Scatterplot showing the relationship between FFMQ Non-reactivity gain (Week 4 − Week 1) and nHF during the post-stress meditation period in the MG. Each point represents one participant, and the line indicates the fitted linear trend.

## Discussion

This study tested whether a brief online mindfulness meditation training alters autonomic regulation during a standardized post-stress meditation sequence. Using a pre/post laboratory protocol combining an acute cognitive stressor (mental arithmetic) and subsequent guided focused-attention breathing meditation, we found a group-specific post-training shift in HRV indices: the meditation group showed lower LF/HF and higher nHF after training, and also differed from an active health-management group in the post-training session. Importantly, this physiological pattern was not merely a generic improvement shared across interventions. Although both groups showed symptom reductions across the four-week period, only the meditation group exhibited evidence linking physiological modulation to mindfulness skill acquisition: within the meditation group, training-related improvement in Non-reactivity was associated with higher nHF during the post-stress meditation period. Together, these findings support the view that brief online mindfulness training can change autonomic responses in the transition from stress to meditation, and that individual differences in acceptance-related skill gains track the degree of post-stress parasympathetic modulation during meditation.

Regarding subjective outcomes, symptom measures decreased toward the later weeks in both groups. Because these improvements were not group-specific, they likely reflect active ingredients present in each condition rather than a mindfulness-specific pathway. In the control group, health-management components (e.g., supplementation and self-monitoring via a food diary) may have contributed to symptom changes, consistent with prior reports that nutritional interventions can influence mood- and anxiety-related outcomes (e.g., ^[16–19]^). Accordingly, symptom measures alone provided limited leverage for isolating mindfulness-specific effects in this brief intervention.

Importantly, mindfulness-related self-report showed a group-specific change: the FFMQ Observing facet increased over the training period in the MG but not in the CG. The FFMQ is one of the most widely used self-report measures of dispositional mindfulness, and its construct validity has been supported across meditating and non-meditating samples, including in active-controlled trial contexts^[14,15]^. The Observing facet captures the tendency to notice and attend to internal and external experiences^[14]^. Moreover, FFMQ scores have been reported to change during mindfulness-base training, including increases across the training period^[15,20]^. Practice engagement has also been associated with changes in FFMQ scores^[21,22]^. Therefore, the MG-specific increase in Observing is consistent with the interpretation that the online mindfulness training strengthened mindful attention/awareness at the subjective level.

Furthermore, physiological outcomes also showed a group-specific pattern in HRV after training. In the post-training session, MG showed lower LF/HF and higher nHF than CG, and within MG these indices shifted from pre-to post-training (LF/HF decreased; nHF increased). Together, these results indicate that the training period was accompanied by a systematic change in frequency-domain HRV indices that was specific to MG.

LF/HF and HF-derived indices are widely used frequency-domain HRV measures^[8]^. LF/HF has often been discussed as an index of sympathetic predominance or “sympathovagal balance,” but its physiological specificity is debated; thus, LF/HF should be interpreted cautiously^[23–26]^. We therefore emphasize HF-related indices, including nHF, which are more commonly discussed as reflecting vagal cardiac modulation, while reporting LF/HF as a supplementary measure^[8,9]^. With these caveats in mind, the MG-specific post-training shift—higher nHF and lower LF/HF—is consistent with increased parasympathetic modulation during the post-stress meditation sequence and aligns with reports that mindfulness or breath-focused guided practices can be accompanied by increases in vagally mediated HRV indices (e.g.,^[27,28]^).

Several factors may contribute to this pattern. Mindfulness training may enhance attentional stability on an interoceptive anchor and reduce perseverative cognition such as rumination—mechanisms emphasized in contemporary accounts of mindfulness training^[13]^ and linked to autonomic regulation and physiological load in prior work^[29,30]^. In addition, because the session included a guided focused-attention breathing meditation, the post-training HRV pattern may partly reflect more effective implementation of breath-based regulation during the session, consistent with reports that focused-attention meditation can facilitate parasympathetic cardiac modulation^[28]^. At the same time, a conservative interpretation is warranted because HF-related indices (including nHF) are sensitive to respiration, and methodological guidance notes that respiratory changes can substantially influence HF power and related metrics^[8,9,25]^. Because respiration was not measured, we cannot disentangle training-related changes in autonomic regulation from breathing-driven contributions during the guided meditation; mechanistic claims should therefore be treated as provisional.

Nevertheless, we observed evidence linking the HRV pattern to mindfulness-specific skill acquisition, beyond a purely breathing-driven account. Within MG, the only robust psychophysiological association was a positive correlation between training-related improvement in FFMQ Non-reactivity and nHF during the meditation period in the post-training session. This relationship provides a plausible bridge between skill acquisition and physiology: individuals who reported a greater tendency to experience internal events without automatic reaction also showed stronger vagal modulation while practicing meditation. Because Non-reactivity reflects an acceptance-oriented stance^[14]^ rather than breathing mechanics per se, this convergence lends additional support to the interpretation that the observed HRV shift is meaningfully related to mindfulness training. Conceptually, acceptance is emphasized as a key ingredient through which mindfulness training translates monitoring skills into stress-relevant benefits^[13]^.

Several limitations should be considered. First, the sample consisted of healthy young adults with relatively low baseline symptoms, which may limit generalizability and reduce the likelihood of detecting group differences in self-report measures. Second, respiratory rate was not measured; because HF-related indices are sensitive to respiration, future studies should monitor breathing and/or include paced-breathing controls to quantify breathing-related contributions to HRV. Finally, although we used robust non-parametric factorial analysis (ART-ANOVA) and controlled multiple comparisons where appropriate, replication with larger samples would improve precision and enable more definitive conclusions using multiple HRV metrics (e.g., time-domain measures) and, ideally, additional biomarkers of stress physiology. Despite these limitations, the present findings provide a basis for evaluating short online mindfulness training with a combined subjective–physiological approach.

## Conclusion

This study demonstrates that a brief, four-week online mindfulness meditation training produces group-specific shifts in frequency-domain HRV during a standardized post-stress meditation sequence. While both groups showed reductions in symptom measures over time, the meditation group exhibited increased mindfulness-related skills together with HRV changes consistent with greater parasympathetic modulation (higher nHF and lower LF/HF) in the post-training session. Moreover, within the meditation group, training-related improvement in Non-reactivity covaried with nHF during the post-training meditation period. These convergent findings support the utility of combining self-report and psychophysiological measures to test how skill acquisition relates to autonomic regulation during post-stress meditation.

## Materials & methods

### Participants

Forty-three healthy volunteers without any previous mindfulness experience or any other type of meditation practice participated in our study. Three participants were excluded because of recording issues. The final data set included forty participants (17 females, mean age ± std = 21.2 ± 1.39 years) Participants were randomly assigned to one of two groups. These groups were trained to reduce mental distress, such as anxiety and depression. One group performed the training of mindfulness meditation four consecutive weeks (meditation group: MG, n = 20, eight females, 21.2 ± 1.50) or to another group in which they performed the health management program four consecutive weeks (control group: CG, n = 20, nine females, 21.3 ± 1.30). All participants gave written informed consent to participate in the study. After the completion of the study, participants were compensated for their time. The experiments were approved by the ethics committee of the School of Science and Technology, Meiji University, and were conducted according to the principles and guidelines of the Declaration of Helsinki.

### Procedure

This experiment consisted of three phases: pre-training phase, training phase, and post-training phase (Figure1). Before the training phase, the level of mindfulness, depression, anxiety, perceived stress, and personality of each participant was examined using the following three psychological scales: Japanese version of Five Facet Mindfulness Questionnaire (FFMQ^[31]^), the Japanese version of the Beck Depression Inventory-Second Edition (BDI-II^[32]^), the Japanese version of State-Trait Anxiety Inventory (STAI)^[33,34]^.

Pre- and post-training phase were composed as follow: pre-rest period followed mental arithmetic task, meditation, and post-rest period. During each rest period participants were instructed to rest, without engaging in any specific task or mental activity. Each rest period lasted for five minutes. In the mental arithmetic task^[35]^, the participants were required to serially subtract the number 13 from 1,022 as fast and as accurately as possible. Participants were required to answer by pressing the number key with the index fingers of their right hand. They had to restart at 1,022 once they made an error. This task lasted for 10 minutes. During the meditation period, which was based on focused attention (FA) on the breath, participants were given an available guided audio with FA instructions (“breath meditation”, Japanese translation^[36]^). This meditation lasted for seven minutes. During both resting and meditation participants were enjoined to keep their eyes closed. Participants filled out the Japanese version of STAI after the pre-rest, mental arithmetic task, and meditation period.

Training phase began the day after the pre-training phase. In training phase, participants participated in different mental distress reduction training program between groups. MG participated in a four-week online meditation program and practiced each week’s meditation following the audio guidance every day. CG participated in a four-week health management program and consumed nutrients effective for mental distress reduction and kept a food diary. Weekly meeting was held to review the previous week and outline the next week’s theme. Participants filled out FFMQ, BDI-II, STAI every weekend.

### Recordings

ECG signals were recorded from lead II using wireless bio-signal recording device (Polymate Pocket MP208, Miyuki Giken, Tokyo, Japan). An electrode placed on the outer right ankle served as a reference. Electrical activity was amplified and digitized at 1000 Hz.

### ECG data analysis

ECG data were processed using MATLAB (R2024a, MathWorks Inc, Natick, MA, USA). To calculate instantaneous heart rate (HR) in beats per minute (bpm), we detected the interval between R-waves, the most prominent peaks of the ECG. Three participants were excluded from the analysis because the peaks could not be detected due to excessive motion noises (MG: n = 2, CG: n = 1). To measure heart rate variability (HRV), the time series of instant heart rate were calculated using linear interpolation. Power spectral analysis for HRV was performed by the wavelet transforms were applied to the interpolated instant heart rate series to identify frequency power. We calculated power of the low-frequency (LF) (0.04–0.15 Hz) band and high-frequency (HF) (0.15–0.40 Hz) band, and derived LF/HF and normalized HF power (nHF = HF/(LF+HF)). LF/HF and nHF were used as frequency-domain HRV indices in subsequent analyses. The higher the LF/HF value, the more sympathetic activity is dominant, and the lower the value, the more parasympathetic activity is dominant. The LF/HF calculated the power ratio of the LF to HF band:

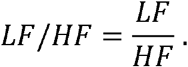

The normalized HF (nHF) is considered as an index of parasympathetic activity^[37]^, calculated the power ratio of the HF to sum of LF and HF:

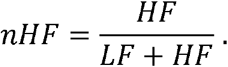

### Psychological scales

In this study, the following three psychological scales were given for all participants.

The Japanese version of FFMQ^[14]^, translated by Sugiura et al.^[31]^, is a 39-item constructed to assess five facets of mindfulness, including Observing, Describing, Acting with Awareness, Nonjudging of Experience, and Non-reactivity to Experience. The measure uses a five-point Likert scale, ranging from 1 (never or rarely true) to 5 (very often or always true).

The Japanese version of BDI-II^[38]^, translated by Kojima et al.^[32]^, is 21-item constructed to measure the severity of depressive symptoms. The measure uses a four-point Likert scale, ranging from 0 to 3 (e.g., “I do not feel sad”; “I feel sad much of the time”; “I am sad all the time”; “I am so sad and unhappy I can’t stand it”).

The Japanese version of STAI^[33]^, translated by Shimizu & Imae^[34]^, is 20-item constructed to measure the state-anxiety (10-item) and trait-anxiety (10-item). The measure uses a four-point Likert scale, ranging from 1 (strongly disagree) to 4 (strongly agree).

### Statistical analysis

The scores of psychological scales, LF/HF, and nHF data were first rank transformed, a robust technique for analyzing non-parametric factorial data using the Align Rank Transform (ART) tool (for details, see^[39]^). The ART method enables the analysis of interaction effects and main effects using standard ANOVA procedures—an approach typically unavailable with conventional non-parametric tests^[39,40]^. This technique is particularly advantageous when data exhibit significant deviations from normality, as it preserves statistical power while minimizing the risk of Type I errors^[40]^. Additionally, the ART procedure has demonstrated statistical power comparable to, or even exceeding, the parametric F-test, particularly under conditions where the assumptions of normal theory are violated^[41]^.

First, to evaluate the effect of training on score of psychological scales (FFMQ, BDI-II, STAI), a two-way ANOVA with ART was conducted with Group (MG or CG) as a between-subject factor and Week (pre-screening, week 1, week 2, week 3, or week 4) as a within-subject factor. Second, to evaluate the effect of training on the STAI state in each period, a three-way ANOVA with ART was performed with Group (MG or CG) as the between-subjects factor, Phase (pre-training or post-training) and Period (after pre-Rest, after stress task or after post-Rest) as the within-subject factors. Third, to identify change rate of autonomic system activities from pre-rest period, the power ratio of the pre-rest period to other periods (mental arithmetic task, meditation, and post-rest period) was calculated. The change ratio of LF/HF and nHF were conducted a three-way ANOVA with ART was performed with Group (MG or CG) as the between-subjects factor, Phase (pre-training or post-training) and Period (pre rest, stress task, or post rest) as the within-subject factors. Finally, to find relations between effect of meditation training and autonomic system activities after meditation training, the Spearman’s rank correlation coefficient was calculated between the FFMQ score at week 1 subtracted from that at week 4 and the nHF or LF/HF for each period after training (Bonferroni corrected: *p* = 0.05/27 (3 periods × 9 psychological scales).

## Author Contributions

Yuki Tsuji (Methodology, Formal analysis, Visualization, Writing-original draft), Itsuki Kondo (Investigation), and Sotaro Shimada (Conceptualization, Supervision, Writing-original draft, Writing-review&editing).

## Data availability

The data that support the findings of this study are available from the corresponding author upon reasonable request.

## References

1. Kabat-Zinn, J. Mindfulness-based interventions in context: past, present, and future. Clin. Psychol. Sci. Pract. 10, 144–156 (2003).

2. Sommers-Spijkerman, M., Austin, J., Bohlmeijer, E. & Pots, W. New Evidence in the Booming Field of Online Mindfulness: An Updated Meta-analysis of Randomized Controlled Trials. JMIR Ment Health 8, e28168 (2021).

3. Witarto, B. S. et al. Effectiveness of online mindfulness-based interventions in improving mental health during the COVID-19 pandemic: A systematic review and meta-analysis of randomized controlled trials. PLoS ONE 17, e0274177 (2022).

4. Reangsing, C., Trakooltorwong, P., Maneekunwong, K., Thepsaw, J. & Oerther, S. Effects of online mindfulness-based interventions (MBIs) on anxiety symptoms in adults: a systematic review and meta-analysis. BMC Complement Med Ther 23, 269 (2023).

5. Creswell, J. D. & Goldberg, S. B. The meditation app revolution. American Psychologist https://doi.org/10.1037/amp0001576 (2025) doi:10.1037/amp0001576.

6. Macrynikola, N. et al. The impact of mindfulness apps on psychological processes of change: a systematic review. npj Mental Health Res 3, 14 (2024).

7. Abeysinghe Mudiyanselage, C. A. K. R., Ewens, B., Smyth, A., Dickson, J. & Ang, S. G. M. Enablers and Barriers of Online Mindfulness-Based Interventions for Informal Carers: A Mixed-Methods Systematic Review. Mindfulness 15, 1257–1274 (2024).

8. Task Force of the European Society of Cardiology and the North American Society of Pacing and Electrophysiology. Heart Rate Variability: Standards of Measurement, Physiological Interpretation, and Clinical Use. Circulation. 93, 1043–1065 (1996).

9. Laborde, S., Mosley, E. & Thayer, J. F. Heart Rate Variability and Cardiac Vagal Tone in Psychophysiological Research – Recommendations for Experiment Planning, Data Analysis, and Data Reporting. Front. Psychol. 08, (2017).

10. Beuchel, P. & Cramer, C. Heart Rate Variability and Perceived Stress in Teacher Training: Facing the Reality Shock With Mindfulness? Global Advances in Integrative Medicine and Health 12, 27536130231176538 (2023).

11. Kirk, U. & Axelsen, J. L. Heart rate variability is enhanced during mindfulness practice: A randomized controlled trial involving a 10-day online-based mindfulness intervention. PLoS ONE 15, e0243488 (2020).

12. Kirk, U. et al. App-Based Mindfulness for Attenuation of Subjective and Physiological Stress Reactivity in a Population With Elevated Stress: Randomized Controlled Trial. JMIR Mhealth Uhealth 11, e47371 (2023).

13. Lindsay, E. K. & Creswell, J. D. Mechanisms of mindfulness training: Monitor and Acceptance Theory (MAT). Clinical Psychology Review 51, 48–59 (2017).

14. Baer, R. A. et al. Construct Validity of the Five Facet Mindfulness Questionnaire in Meditating and Nonmeditating Samples. Assessment 15, 329–342 (2008).

15. Goldberg, S. B. et al. Does the Five Facet Mindfulness Questionnaire measure what we think it does? Construct validity evidence from an active controlled randomized clinical trial. Psychological Assessment 28, 1009–1014 (2016).

16. Kikuchi, A. M., Tanabe, A. & Iwahori, Y. A systematic review of the effect of L-tryptophan supplementation on mood and emotional functioning. Journal of Dietary Supplements 18, 316–333 (2021).

17. Oliveira, I. J. L. D., De Souza, V. V., Motta, V. & Da-Silva, S. L. Effects of Oral Vitamin C Supplementation on Anxiety in Students: A Double-Blind, Randomized, Placebo-Controlled Trial. Pakistan J. of Biological Sciences 18, 11–18 (2014).

18. Sim, M. et al. Vitamin C supplementation promotes mental vitality in healthy young adults: results from a cross-sectional analysis and a randomized, double-blind, placebo-controlled trial. Eur J Nutr 61, 447–459 (2022).

19. Tao, Y. et al. Impact of Vitamin B1 and Vitamin B2 Supplementation on Anxiety, Stress, and Sleep Quality: A Randomized, Double-Blind, Placebo-Controlled Trial. Nutrients 17, 1821 (2025).

20. Baer, R. A., Carmody, J. & Hunsinger, M. Weekly Change in Mindfulness and Perceived Stress in a Mindfulness□Based Stress Reduction Program. J Clin Psychol 68, 755–765 (2012).

21. Carmody, J. & Baer, R. A. Relationships between mindfulness practice and levels of mindfulness, medical and psychological symptoms and well-being in a mindfulness-based stress reduction program. J Behav Med 31, 23–33 (2008).

22. Goldberg, S. B., Del Re, A. C., Hoyt, W. T. & Davis, J. M. The secret ingredient in mindfulness interventions? A case for practice quality over quantity. Journal of Counseling Psychology 61, 491–497 (2014).

23. Eckberg, D. L. Sympathovagal Balance: A Critical Appraisal. Circulation 96, 3224–3232 (1997).

24. Heathers, J. A. J. Sympathovagal balance from heart rate variability: an obituary. Experimental Physiology 97, 556–556 (2012).

25. Quintana, D. S. & Heathers, J. A. J. Considerations in the assessment of heart rate variability in biobehavioral research. Front. Psychol. 5, (2014).

26. Billman, G. E. The LF/HF ratio does not accurately measure cardiac sympatho-vagal balance. Front. Physio. 4, (2013).

27. Ditto, B., Eclache, M. & Goldman, N. Short-term autonomic and cardiovascular effects of mindfulness body scan meditation. ann. behav. med. 32, 227–234 (2006).

28. Ooishi, Y., Fujino, M., Inoue, V., Nomura, M. & Kitagawa, N. Differential Effects of Focused Attention and Open Monitoring Meditation on Autonomic Cardiac Modulation and Cortisol Secretion. Front. Physiol. 12, 675899 (2021).

29. Ottaviani, C. et al. Physiological concomitants of perseverative cognition: A systematic review and meta-analysis. Psychological Bulletin 142, 231–259 (2016).

30. Cropley, M. et al. The Association between Work-Related Rumination and Heart Rate Variability: A Field Study. Front. Hum. Neurosci. 11, (2017).

31. Sugiura, Y., Sato, A., Ito, Y. & Murakami, H. Development and Validation of the Japanese Version of the Five Facet Mindfulness Questionnaire. Mindfulness 3, 85–94 (2012).

32. Kojima, M. et al. Cross-cultural validation of the Beck Depression Inventory-II in Japan. Psychiatry Research 110, 291–299 (2002).

33. Spielberger, C. D., Gorsuch, R. L., & Lushene. R. E. Manual for the State-Trait Anxiety Inventory. Palo Alto, CA: Consulting Psychologists Press. (Consulting Psychologists Press, Palo Alto, CA, 1970).

34. Shimizu, H. & Imae, K. Development of the Japanese edition of the Spielberger state-trait anxiety inventory (STAI) for student use. The Japanese Journal of Educational Psychology 29, 348–353 (1981).

35. Yao, Z., Zhang, L., Jiang, C., Zhang, K. & Wu, J. Stronger cortisol response to acute psychosocial stress is correlated with larger decrease in temporal sensitivity. PeerJ 4, e2061 (2016).

36. Williams, J. M. G. & Penman, D. Mindfulness: An Eight-Week Plan for Finding Peace in a Frantic World. (Rodale Books, Emmaus, Pa., 2012).

37. Burr, R. L. Interpretation of Normalized Spectral Heart Rate Variability Indices In Sleep Research: A Critical Review. Sleep 30, 913–919 (2007).

38. Beck, A. T., Steer, R. A. & Brown, G. Beck Depression Inventory–II. 10.1037/t00742-000 (2011).

39. Wobbrock, J. O., Findlater, L., Gergle, D. & Higgins, J. J. The aligned rank transform for nonparametric factorial analyses using only anova procedures. in Proceedings of the SIGCHI Conference on Human Factors in Computing Systems 143–146 (ACM, Vancouver BC Canada, 2011). doi:10.1145/1978942.1978963.

40. Durner, E. Effective Analysis of Interactive Effects with Non-Normal Data Using the Aligned Rank Transform, ARTool and SAS® University Edition. Horticulturae 5, 57 (2019).

41. Richter, S. J. Nearly exact tests in factorial experiments using the aligned rank transform. Journal of Applied Statistics 26, 203–217 (1999).

