## Supplementary material for "Brief online mindfulness meditation training modulates heart rate variability during post-stress meditation": Table S1

**Table S1. Results of psychological scales at pre-screening. Mean values of the psychological scales are shown with standard deviation.**

| **Psychological scales** | **Meditation Group**  **(n = 20)** | **Control Group**  **(n = 20)** | ***p* value**  **(group difference)** |
| --- | --- | --- | --- |
| FFMQ total | 110 ± 14.7 | 112 ± 17.7 | 0.753 |
| observing | 23.8 ± 4.73 | 23.3 ± 6.03 | 0.814 |
| describing | 20.8 ± 6.36 | 21.5 ± 8.99 | 0.878 |
| acting with awareness | 23.7 ± 5.89 | 23.8 ± 6.71 | 0.587 |
| nonjudging of experience | 21.0 ± 5.62 | 23.3 ± 6.84 | 0.297 |
| non-reactivity to experience | 21.6 ± 4.12 | 20.5 ± 3.22 | 0.378 |
| STAI-state | 38.7 ± 9.50 | 39.6 ± 11.2 | 0.794 |
| STAI-trait | 47.3 ± 9.02 | 46.5 ± 10.0 | 0.653 |
| BDI-II | 7.70 ± 5.88 | 8.80 ± 6.61 | 0.682 |
