## Supplementary material for "Brief online mindfulness meditation training modulates heart rate variability during post-stress meditation": Table S2

**Table S2. Results of a two-way ANOVA with ART for psychological scales.**

| **Psychological scales** | **Group** | **Week** | **Group × Week** |
| --- | --- | --- | --- |
| FFMQ total | *F*(1,38) = 0.0168 | *F*(4,152) = 2.65 * | *F*(4,152) = 1.24 |
| observing | *F*(1,38) = 0.969 | *F*(4,152) = 3.03 * | *F*(4,152) = 3.33 * |
| describing | *F*(1,38) = 5.10×10^-6^ | *F*(4,152) = 2.59 * | *F*(4,152) = 0.577 |
| acting with awareness | *F*(1,38) = 2.88×10^-3^ | *F*(4,152) = 0.504 | *F*(4,152) = 0.588 |
| nonjudging of experience | *F*(1,38) = 1.00 | *F*(4,152) = 2.31 | *F*(4,152) = 1.85 |
| non-reactivity to experience | *F*(1,38) = 1.59 | *F*(4,152) = 0.712 | *F*(4,152) = 0.0583 |
| STAI-state | *F*(1,38) = 0.385 | *F*(4,152) = 4.04 * | *F*(4,152) = 1.51 |
| STAI-trait | *F*(1,38) = 0.229 | *F*(4,152) = 4.22 * | *F*(4,152) = 0.496 |
| BDI-II | *F*(1,38) = 1.93 | *F*(4,152) = 8.60 * | *F*(4,152) = 0.471 |

Note:* *p* < 0.05
